# Using RNA-targeting CRISPR-Cas13 and engineered U1 systems to reduce *ABCA4* splice variants in Stargardt disease

**DOI:** 10.1101/2024.03.08.584155

**Authors:** Roxanne Hsiang-Chi Liou, Daniel Urrutia-Cabrera, Ida Maria Westin, Irina Golovleva, Guei-Sheung Liu, Satheesh Kumar, Samuel McLenachan, Fred Kuanfu Chen, Fei-Ting Hsu, Tom Edwards, Keith R Martin, Albert Wu Cheng, Raymond C.B. Wong

**Affiliations:** Centre for Eye Research Australia, Royal Victorian Eye and Ear Hospital. Australia; Ophthalmology, Department of Surgery, University of Melbourne, Australia; Medical and Clinical Genetics, Department of Medical Biosciences, University of Umeå, Sweden; Centre for Ophthalmology and Visual Science, University of Western Australia, Australia; Lions Eye Institute, Western Australia, Australia; Department of Biological Science and Technology, China Medical University, Taiwan; Department of Clinical Neurosciences, University of Cambridge, United Kingdom; School of Biological and Health Systems Engineering, Arizona State University, United States; State Key Laboratory of Stem Cell and Reproductive Biology, Institute of Zoology, Chinese Academy of Sciences, Beijing, China; Key Laboratory of Organ Regeneration and Reconstruction, Chinese Academy of Sciences, Beijing, China; Beijing Institute for Stem Cell and Regenerative Medicine, Beijing, China; University of Chinese Academy of Sciences, Beijing, China

## Abstract

Dysregulation of the alternative splicing process results in aberrant mRNA transcripts, leading to dysfunctional proteins or nonsense-mediated decay that cause a wide range of mis-splicing diseases. Development of therapeutic strategies to target the alternative splicing process could potentially shift the mRNA splicing from disease isoforms to a normal isoform and restore functional protein. As a proof of concept, we focus on Stargardt disease (STGD1), an autosomal recessive inherited retinal disease caused by biallelic genetic variants in the *ABCA4* gene. The splicing variants c.5461-10T>C and c.4773+3A>G in *ABCA4* cause the skipping of exon 39-40 and exon 33-34 respectively. In this study, we compared the efficacy of different RNA-targeting systems to modulate these *ABCA4* splicing defects, including four CRISPR-Cas13 systems (CASFx-1, CASFx-3, RBFOX1N-dCas13e-C and RBFOX1N-dPspCas13b-C) as well as an engineered U1 system (ExSpeU1). Using a minigene system containing *ABCA4* variants in the human retinal pigment epithelium ARPE19, our results show that RBFOX1N-dPspCas13b-C is the best performing CRISPR-Cas system, which enabled up to 80% reduction of the mis-spliced *ABCA4* c.5461-10T>C variants and up to 78% reduction of the *ABCA4* c.4773+3A>G variants. In comparison, delivery of a single ExSpeU1 was able to effectively reduce the mis-spliced *ABCA4* c.4773+3A>G variants by up to 84%. We observed that the effectiveness of CRISPR-based and U1 splicing regulation is strongly dependent on the sgRNA/snRNA targeting sequences, highlighting that optimal sgRNA/snRNA designing is crucial for efficient targeting of mis-spliced transcripts. Overall, our study demonstrated the potential of using RNA-targeting CRISPR-Cas technology and engineered U1 to reduce mis-spliced transcripts for *ABCA4*, providing an important step to advance the development of gene therapy to treat STGD1.

## Introduction

Stargardt disease (STGD1) is the most common juvenile macular degeneration, accounting for a significant proportion of blindness in children and young adults. STGD1 is caused by biallelic pathogenic variants in the *ABCA4* gene which encodes a photoreceptor-specific transporter used to facilitate the removal of all-trans-retinal from disc membranes through the visual cycle ^1^. Loss of ABCA4 function results in the accumulation of toxic lipofuscin bisretinoid in the retinal pigment epithelium (RPE), ultimately leading to RPE degeneration and photoreceptor loss ^2^. To date, more than 1200 disease-causing *ABCA4* variants, including missense, splicing, truncating, and frameshift mutations, are associated with STGD1 (HGMD, till January 2023). Previous studies also revealed common intronic variants of the *ABCA4* gene, including the mutations c.5461-10T>C and c.4773+3A>G ^3,4^. Subsequent *in vitro* studies showed that both variants cause aberrant alternative splicing of the *ABCA4* gene; c.5461-10T>C causes skipping of exon 39 and 40 ^5,6^, while c.4773+3A>G causes skipping of exon 33 and 34 ^7,8^.

One of the main challenges for DNA-targeting CRISPR-Cas techniques is the potential off-target effects for genome editing which may lead to permanent adverse effects. In comparison, RNA-targeting CRISPR-Cas systems hold an advantage in the sense that RNA represents a transient entity, as such any potential off-target modification to RNA would not have permanent deleterious effects on the cell. Previous studies have demonstrated the potential of RNA-targeting CRISPR-Cas13 systems as a powerful tool for targeting and regulating alternative splicing processes. For instance, the catalytically inactive form of the Cas13d family member CasRx (RfxCas13d) has been engineered to facilitate the exclusion of exon 10 of *MAPT* gene for treating frontotemporal dementia ^9^. Furthermore, several splicing regulatory domains have been utilized to improve the efficacy of alternative splicing modulation, including RBFOX1 ^10,11^ and RBM38 ^12^. A previous study by Du et al. has reported improved CRISPR-Cas13 splicing modulating system by coupling dCasRx with RBFOX1 at both N- and C-terminal (CASFx-1/RBFOX1N-dCasRx-C) and with RBM38 at the C-terminal (CASFx-3/dCasRx-RBM38) ^13^. The artificially engineered CASFx systems showed better efficacy in inducing exon inclusion compared to dCasRx alone, as dCasRx showed a tendency to cause exon skipping when targeting the branch point or the splice donor and acceptor sites ^9^. In the case of inducing exon inclusion of the *SMN2* gene in spinal muscular atrophy patients, CASFx-1 showed a 21-fold increase while CASFx-3 showed a 6-fold increase in exon inclusion compared to dCasRx ^13^.

Other members of the Cas13 family have also been utilized to target RNA, including Cas13b and Cas13e. Among the Cas13b orthologs, the PspCas13b (derived from *Prevotella sp. P5– 125*) exerted the most prominent efficiency and specificity for RNA knockdown in mammalian cell applications ^14^. The catalytically inactive dPspCas13b has also been engineered into a splicing effector combined with RBFOX1 at both N- and C-terminal (RBFOX1N-dPspCas13b-C) and has shown better efficiency in inducing exon inclusion of the *SMN2* gene compared to CASFx-1 ^13^. In addition, Cas13e is a recently characterized ribonuclease in the Cas13 family and has garnered special interest for its compact size. With a size of 750 aa, Cas13e is one of the smallest Cas proteins in the Cas13 family, making it an attractive target for adeno-associated virus (AAV) packaging. When coupled with engineered deaminase, Cas13e exhibited robust efficiency in RNA base conversion and has demonstrated clinical utility in recent studies ^15^.

Other than CRISPR technologies, engineered U spliceosomal small nuclear RNAs (snRNAs) also serve as promising tools for modulating alternative splicing. During the alternative splicing process, the 5’ splice site is recognised through base-pairing the U1 snRNAs and the intron sequences of the pre-mRNA. When alternative splicing occurs, different splice sites within the pre-mRNA may compete for recognition by the spliceosome. Thus, U1 snRNAs contribute to the selection of specific splice sites, influencing whether an exon is included or excluded in the mature mRNA ^16^. As such, engineered U1 snRNAs have emerged as effective tools for targeted recognition to correct splicing defects. Previous studies have showcased the versatility of engineered U1 snRNAs, demonstrating their capability to induce both inclusion and exclusion of targeted exons in mammalian cells ^17,18^. Recent development of Exon-Specific U1 snRNAs (ExSpeU1) allowed targeting of the non-conserved intronic sequences downstream of the 5′ splice site of the skipped exon. This design lowers the potential of off-target effects and has shown great potential in therapeutic applications ^19,20^. Several studies highlight the therapeutic potential of ExSpeU1 treatment correct splicing defects; ExSpeU1s have been shown to induce exon inclusion for the *coagulation factor VII* (*F7*) gene ^21^, the *coagulation factor IX* (*F9*) gene and *SMN2* gene ^22^ in mammalian cells. Apart from exon inclusion, U1 snRNA treatment also induces skipping of mutated exon 9a of the *RPGR* gene in HEK293, 661W, and PC-12 cell lines ^23^, offering an exon skipping-based approach to correct splicing defects for X-linked retinitis pigmentosa. ExSpeU1s were also tested *in vivo* and exhibited the potential of rescuing alternative splicing defects. *F7* expression and protein level were restored after AAV injection of the ExSpeU1 in mice ^24^. Successful exon inclusion of the *SMN2* gene had also been observed after AAV-mediated delivery of ExSpeU1 in mice ^25^. These advancements underscore the promising role of engineered U1 snRNAs in manipulating alternative splicing and hold great potential for therapeutic interventions in diseases associated with splicing dysregulation.

In this study, we tested the efficacy of several RNA-targeting CRISPR-Cas13 and ExSpeU1 systems as splicing effectors for exon inclusion of *ABCA4* c.4773+3A>G and c.5461-10T>C variants, including ExSpeU1, CASFx-1, CASFx-3 and RBFOX1N-dPspCas13b-C. We also engineered a new Cas13e-based splicing effector by combining dCas13e with RBFOX1 (RBFOX1N-dCas13e-C). Using a minigene system containing *ABCA4* variants in ARPE19 cells, our results highlighted that RBFOX1N-dPspCas13b-C can achieve up to 80% reduction of mis-spliced transcripts for both *ABCA4* c.4773+3A>G and c.5461-10T>C variants. Moreover, ExSpeU1s can reduce *ABCA4* c.4773+3A>G mis-spliced transcripts by up to 84%. These results show that RNA-targeting CRISPR-Cas systems and ExSpeU1s can be designed to manipulate alternative splicing of *ABCA4*, providing a novel approach to mitigate adverse effects of the splicing mutations for treating STGD1.

## Results

### Protein structure prediction of ABCA4 c.5461-10T>C variants with deletion of exon 39-40

The *ABCA4* c.5461-10T>C variant causes the deletion of exons 39 and 40, leading to a premature stop codon within the reading frame. To verify how the variant affects the structure and folding of the ABCA4 protein, we performed *in silico* analysis to predict the protein structure for the wild-type (WT) ABCA4, the ABCA4 Δ exon 39 mutant and the Δ exon 39-40 mutant protein sequences. We utilized ColabFold to predict the three-dimensional protein structures based on the sequences, which generates a faster prediction of protein structures by combining the fast homology search of MMseq2 (Many-against-Many sequence searching) with AlphaFold2 ^26^. Our results revealed structural differences between the WT ABCA4 (Fig. 1A), and the mis-spliced variants (Fig. 1B & C). We further carried out the protein structure comparison of the mis-splice variants with a published human WT ABCA4 experimental structure ^27^. Similarly, the structural comparison revealed aberrant folding of the mutant proteins compared to the WT ABCA4 (Fig. 1D & E), resulting in dysfunctional ABCA4 disease isoforms with a truncated transmembrane domain encoded by exon 39-40.

**Figure 1.**
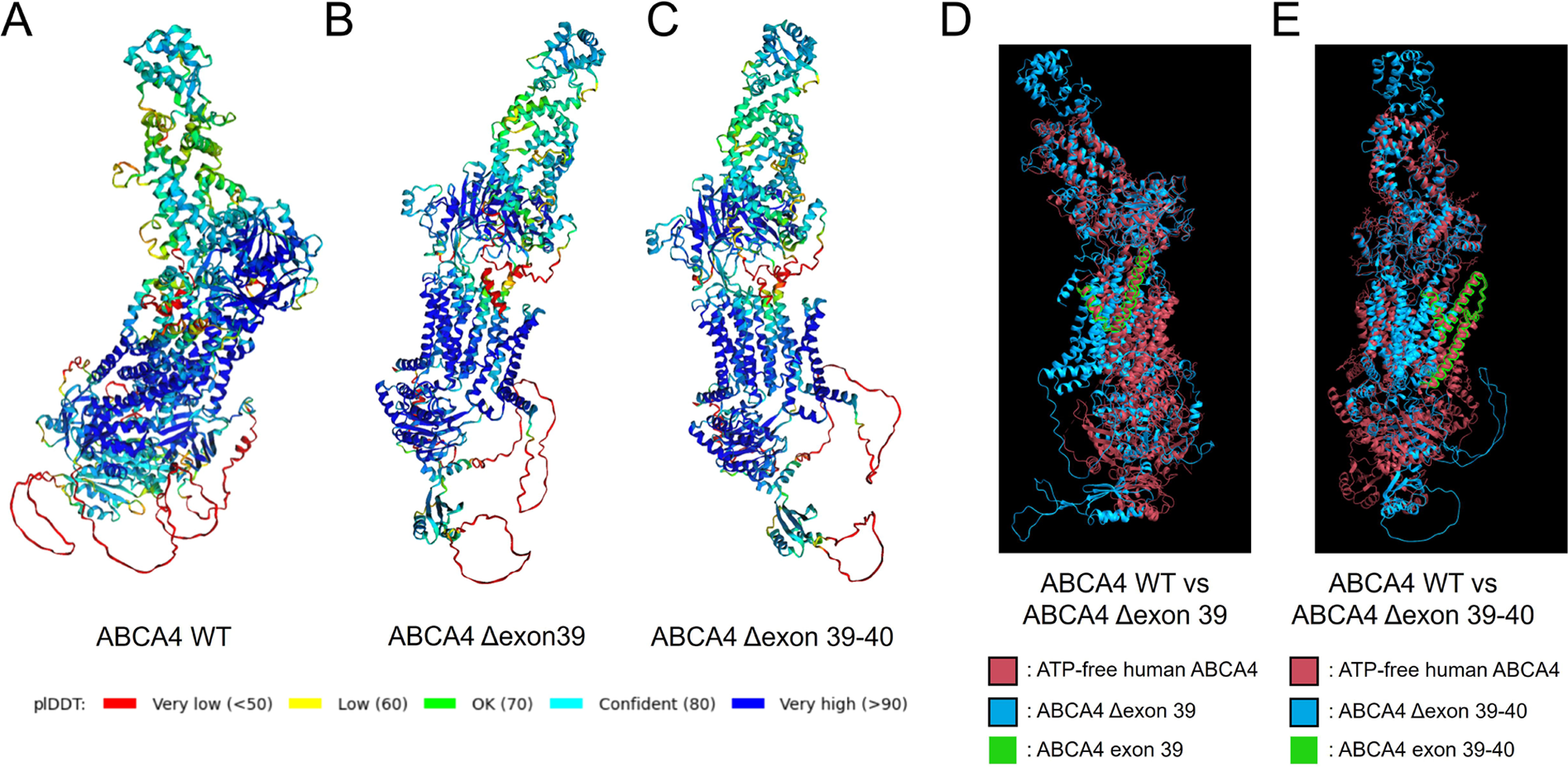
Structural prediction and comparison of the ABCA4 variants with exon 39-40 deletion. (A) Protein structures of the wildtype ABCA4, (B) the ABCA4 Δexon 39, and (C) the ABCA4 Δexon 39-40 predicted by the ColabFold tool. The amino acid residues were coloured based on the plDDT score, with a higher score representing higher confidence in the accuracy of the predicted structure. Regions below 50 plDDT may be unstructured in isolation. (D) Comparison of protein structures of ABCA4 Δexon 39 and WT using ChimeraX. The verified ATP-free human ABCA4 protein structure was used as a reference (red, ^27^). ABCA4 Δexon 39 was presented in blue. The peptide encoded by exon 39 was highlighted in green. (E) Comparison of protein structures of ABCA4 Δexon 39-40 and WT using ChimeraX. The verified ATP-free human ABCA4 protein structure was used as a reference and highlighted in red. ABCA4 Δexon 39-40 was presented in blue. The peptide encoded by exon 39-40 was highlighted in green.

### Evaluation of CASFx to modulate splicing defects of ABCA4 c.5461-10T>C

In order to test the potential of RNA-targeting CRISPR-Cas13 systems to facilitate exon inclusion of the mis-spliced *ABCA4* gene, we utilized an *ABCA4* c.5461-10T>C minigene system containing *ABCA4* exon 39 and 40 along with the c.5461-10T>C variant in the pSPL3 exon-trapping vector ^7^. We first tested the CASFx systems for the inclusion of exon 39 and 40. The dCasRx-based CRISPR splicing factors, CASFx-1 (RBFOX1N-dCasRx-C) and CASFx-3 (dCasRx-RBM38) were tested using the minigene system in the human RPE cell line ARPE19. We designed 2 sgRNAs each for targeting the downstream introns of the target exon 39 or 40 (Fig. 2A). The mutant *ABCA4* minigene (MT) showed three distinctive splicing products after transfection into ARPE19 cells; a full-length (FL) product containing both exon 39 and 40 (514 bp), a truncated *ABCA4* disease isoform with exon 39 deletion (Δ exon 39, 390 bp), and a truncated *ABCA4* disease isoform with deletion of exon 39 and 40 (Δ exon 39-40, 260 bp) (Fig. 2B & C).

**Figure 2.**
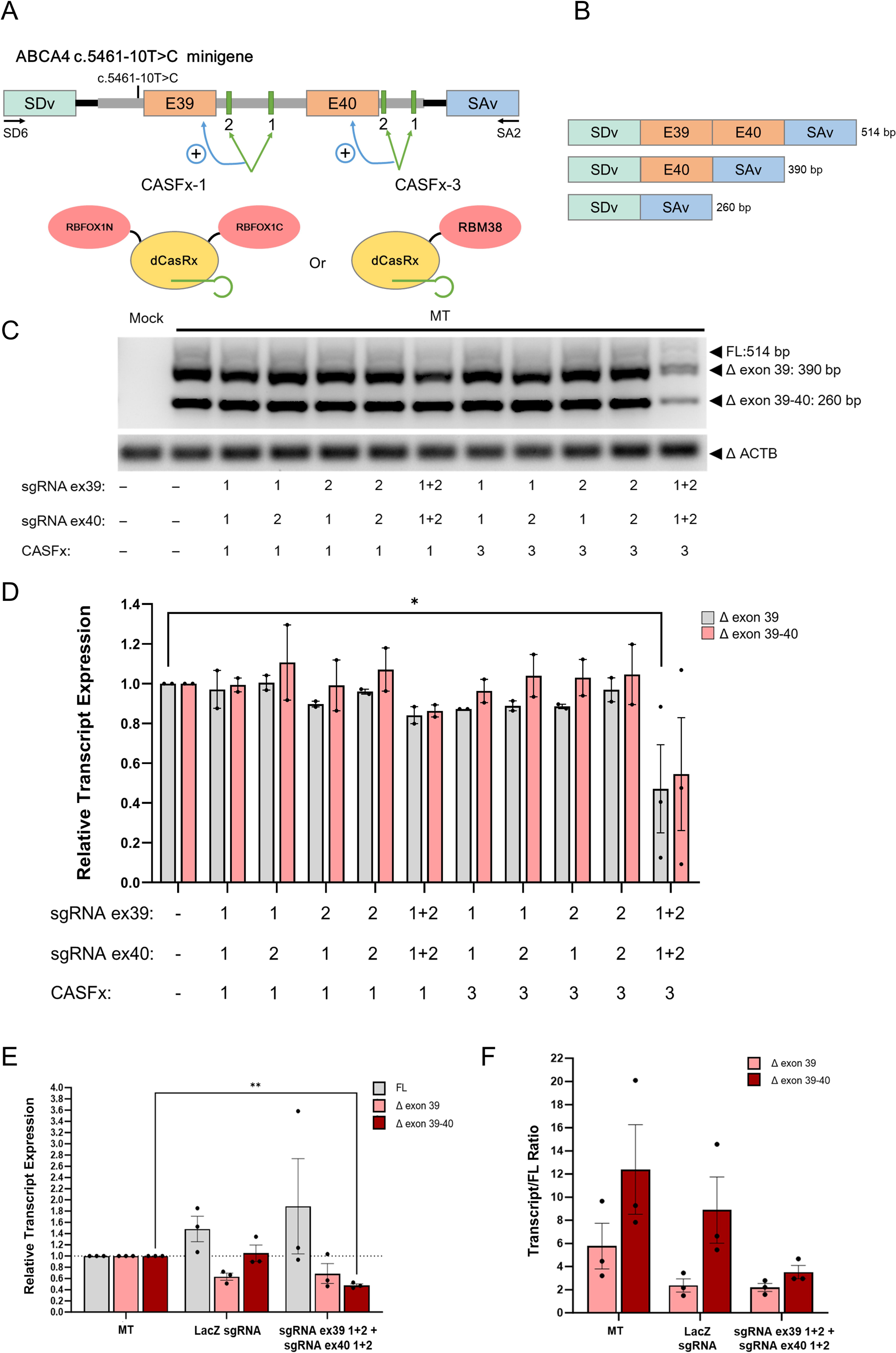
Evaluation of CASFx to modulate splicing defects of ABCA4 c.5461-10T>C. (A) A schematic illustration of the pSPL3–ABCA4 c.5461-10T>C minigene and the CRISPR splicing factors CASFx-1 (RBFOX1N-dCasRx-C) and CASFx-3 (dCasRx-RBM38), as well as two sgRNA target sites downstream of exon 39 (E39) and exon 40 (E40) respectively (green). *ABCA4* exons (orange), and flanking introns (grey) were cloned into the pSPL3 vector (black) along with the mutation between two pSPL3 exons; splice donor vector (SDv, green) and splice acceptor vector (SAv, blue). The *ABCA4* splice variants were amplified using the SD6 and SA2 primers. (B) Expected splicing products from the ABCA4 c.5461-10T>C minigene. Normal splicing of the minigene gives rise to a FL product containing both exon 39 and 40 (514 bp) while skipping of exon 39 or exon 39 and 40 generates shorter isoforms (390 and 260 bp). (C) RT-PCR results of *ABCA4* splice variants in ARPE19 cells transfected with the ABCA4 minigene system (MT), CASFx-1, CASFx-3 and specified sgRNAs. *ACTB* was used as a housekeeping gene. Mock control represents ARPE19 without transfection of the *ABCA4* minigene system. (D) Gel quantification of relative transcript ratio of ABCA4 splice variants compared to MT control without treatment. Data are represented as mean ± SEM (n = 3 for CASFx-3 ex39 sgRNA1+2 ex40 sgRNA 1+2, n = 2 for the rest). *: p < 0.05. (E) qPCR quantification of the relative expression level of *ABCA4* splice variants and (F) relative expression level of disease transcript to FL transcript compared to MT control without treatment. Data are presented as mean ± SEM (n = 3). **: p < 0.01.

We first utilized RT-PCR to screen a range of sgRNA/snRNA for CASFx-1 and CASFx-3. Our RT-PCR results indicated that CASFx-3 with the combination of 4 sgRNAs (2 sgRNAs each for targeting exon 39 and 40) resulted in a significant decrease in the Δ exon 39 mis-spliced transcript (52.9% reduction), and a trend of reduction in the Δ exon 39-40 mis-spliced transcript (45.5% reduction), albeit not statistically significant (Fig. 2C & D). All other sgRNA combinations for CASFx-1 and CASFx-3 showed no significant effect in lowering the level of Δ exon 39 and Δ exon 39-40 mis-spliced transcript (Fig. 2C & D).

To further validate the decrease in disease transcript levels shown with CASFx-3 treatment with the combination of 4 sgRNAs, we performed qPCR analysis to quantify the expression level of each transcript. We also included a control sgRNA targeting a region of the LacZ gene (LacZ sgRNA) to ensure the transfected plasmid ratios were even and that the splicing modulating effect is specific to the *ABCA4* sgRNAs. qPCR analysis showed that CASFx-3 treatment with the combination of 4 sgRNAs resulted in a significant decrease in the Δ exon 39-40 mis-spliced transcript (52.3% reduction), and a trend of reduction in the Δ exon 39 mis-spliced transcript (31.3% reduction), albeit not statistically significant (Fig. 2E). In contrast, qPCR analysis showed no decrease in FL transcript after CASFx-3 treatment with *ABCA4* sgRNAs. We also observed a trend of decrease in the Δ exon 39 disease isoform in LacZ sgRNA treated groups, although not statistically significant.

Next, we evaluated the relative expression levels of the disease transcripts to FL by qPCR. Quantification of the disease transcript/FL expression level showed a trend of decrease for both disease transcripts in *ABCA4* sgRNAs-treated group compared to the control (ΔE39/FL = 0.38 fold, ΔE39-40/FL = 0.28 fold compared to control, Fig. 2F), indicating a decrease in the expression of disease transcripts while the expression of FL transcript is maintained. However, the decrease did not reach statistical significance. LacZ sgRNA-treated group also showed a statistically insignificant decrease in disease transcript/FL expression level for both Δ exon 39 and Δ exon 39-40 mis-spliced transcript. (Fig. 2F). Altogether, our results showed that CASFx-3 coupled with multi-sgRNA approach can reduce disease variants of ABCA4 by up to 52%.

### Evaluation of dCas13e to modulate splicing defects of ABCA4 c.5461-10T>C

In order to improve the efficiency of exon inclusion of exon 39 and 40, we utilized another CRISPR-Cas13 system, dCas13e, for alternative splicing modulation. The engineered dCas13e is one of the smallest Cas13 proteins, holding great value in clinical translation by packaging into AAVs. We created a multiplex vector system that expresses both sgRNA and dCas13e coupled with the splicing factor RBFOX1 at both N- and C-terminal in a single all-in-one plasmid (RBFOX1N-dCas13e-C, Fig. 3A). Our RT-PCR results showed that the negative control, a mCherry targeting sgRNA, showed no effect on the level of full-length and disease transcripts as expected. However, the *ABCA4* sgRNA treatment also exerted no significant effect on the disease transcript levels (Δ exon 39 = 0.78 fold, Δ exon 39-40 = 0.76 fold compared to control), as well as the disease transcript/FL ratio (Δ E39/FL = 1.04 fold; Δ E39-40/FL = 0.96 fold, Fig. 3B-D). Collectively, we did not observe a significant change in *ABCA4* disease isoforms using the RBFOX1N-dCas13e-C system.

**Figure 3.**
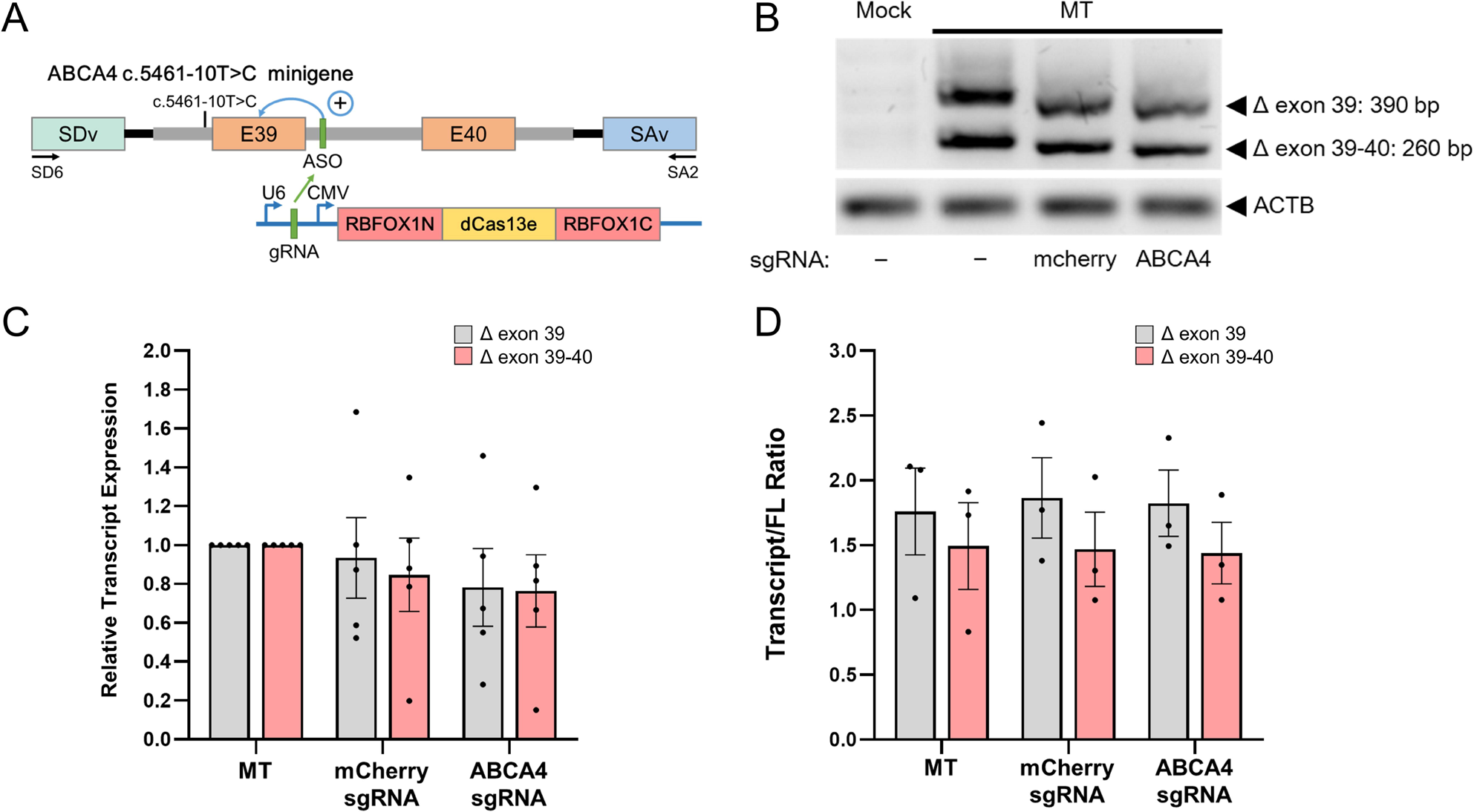
Evaluation of dCas13e to modulate splicing defects of ABCA4 c.5461-10T>C. (A) A schematic illustration of the pSPL3–*ABCA4* c.5461-10T>C minigene and the all-in-one construct of a U6 promoter-driven sgRNA sequence followed by the CMV promoter-driven RBFOX1N-dCas13e-C factor. The sgRNA targeted the same region downstream of E39 (green) as a previously reported antisense oligonucleotide target site. Positions of SD6 and SA2 primers for splicing product amplification are displayed. (B) RT-PCR results of ABCA4 splice variants in ARPE19 cells transfected with the ABCA4 minigene system (MT), RBFOX1N-dCas13e-C, sgRNAs targeting mCherry (control) or *ABCA4*. *ACTB* was used as a housekeeping gene. Mock control represents ARPE19 without transfection of the ABCA4 minigene system. (C) Quantification of relative transcript ratio of *ABCA4* splice variants compared to MT control without treatment. Data are represented as mean ± SEM (n = 5). (D) Quantification of the ratio of disease transcript to FL transcript compared to MT control without treatment. Data are represented as mean ± SEM (n = 3).

### Evaluation of dPspCas13b to modulate splicing defects of ABCA4 c.5461-10T>C

Next, we tested the RBFOX1N-dPspCas13b-C system for splicing modulation of the *ABCA4* c.5461-10T>C splicing variant. 2 sgRNAs were designed to target the mutation region upstream of exon 39 (Fig. 4A). Notably based on RT-PCR results, both sgRNA 1 and sgRNA 2 using the RBFOX1N-dPspCas13b-C system were able to significantly reduce the levels of Δ exon 39 and Δ exon 39-40 disease isoforms in ARPE19 (Fig. 4B). sgRNA 1 showed a significant decrease in Δ exon 39 isoform and Δ exon 39-40 isoform (57.9% reduction and 53.8% reduction respectively). Similarly, sgRNA 2 decreased the Δ exon 39 isoform significantly by 54.2% and Δ exon 39-40 isoform by 51.5% (Fig. 4C).

**Figure 4.**
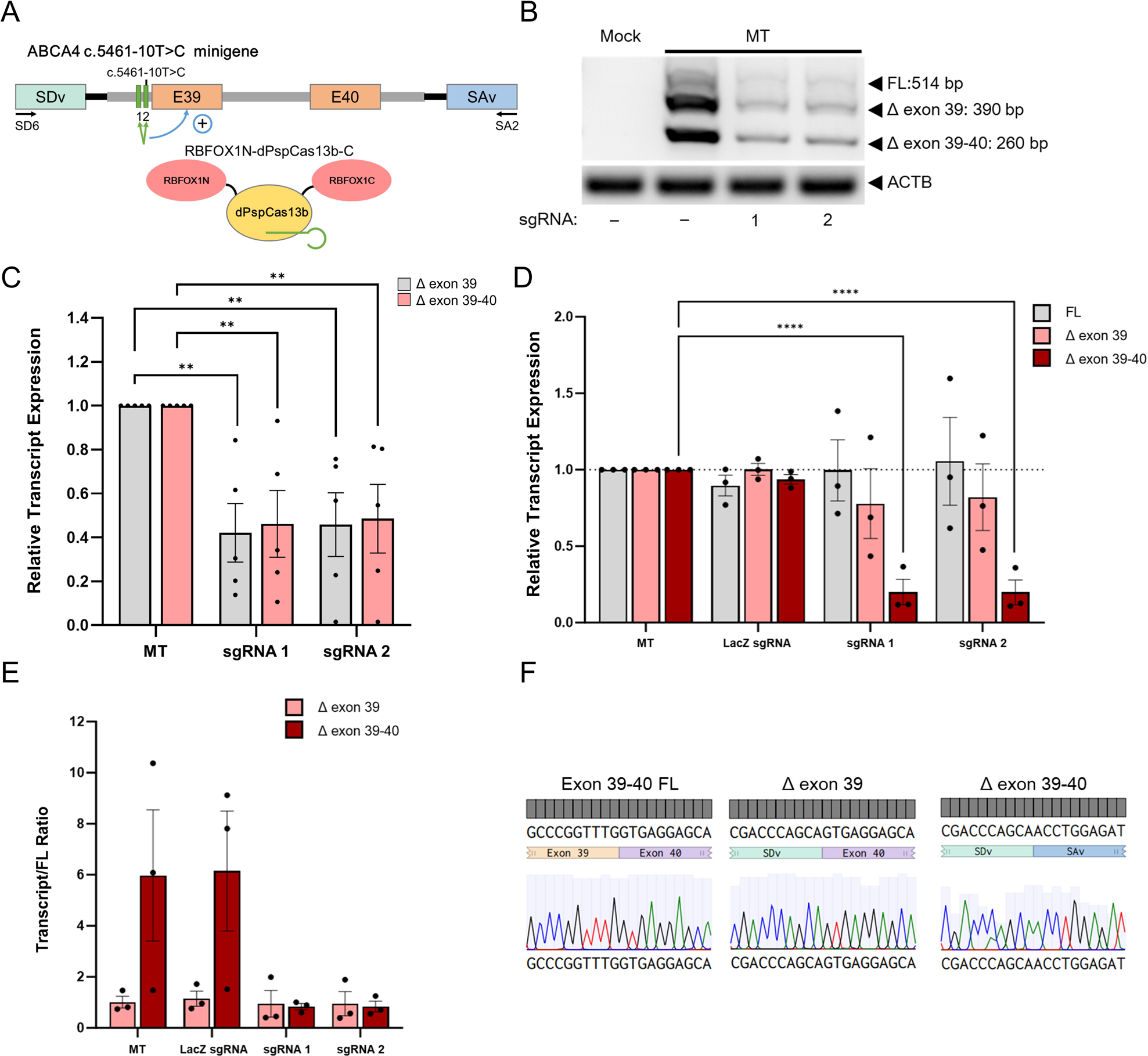
Evaluation of dPspCas13b to modulate splicing defects of ABCA4 c.5461-10T>C. (A) A schematic illustration of the pSPL3–ABCA4 c.5461-10T>C minigene and the CRISPR splicing factor RBFOX1N-dPspCas13b-C, as well as two sgRNA sites targeting the mutation upstream of E39 (green). Positions of SD6 and SA2 primers for splicing product amplification are displayed. (B) RT-PCR results of ABCA4 splice variants in ARPE19 cells transfected with the ABCA4 minigene system (MT), RBFOX1N-dPspCas13b-C and specified sgRNAs. ACTB was used as the loading control. Mock control represents ARPE19 without transfection of the ABCA4 minigene system. (C) Quantification of relative transcript ratio of ABCA4 splice variants compared to MT control without treatment. Data are represented as mean ± SEM (n = 5). **: p < 0.01. (D) qPCR quantification of the relative expression level of ABCA4 splice variants and (E) relative expression level of disease transcript to FL transcript compared to MT control without treatment. Data are presented as mean ± SEM (n = 3). ****: p < 0.0001. (F) Sanger sequencing confirmed the presence of ABCA4 FL transcript (left), the Δexon 39 isoform (middle) and the Δexon 39-40 isoform (right).

Further qPCR validation showed a significant decrease in the expression level of Δ exon 39-40 disease isoforms for both sgRNA 1 (79.9% reduction) and sgRNA 2 (80.1% reduction). Both sgRNA 1 and sgRNA 2 caused a slight but not significant decrease in Δ exon 39 disease isoforms (sgRNA 1 = 0.78 fold, sgRNA 2 = 0.82 fold compared to control). LacZ sgRNA treatment showed no effect on both disease and FL isoforms (Fig. 4D). For the disease transcript/FL expression level, sgRNA 1 caused a sharp reduction of Δ exon 39-40/FL ratio (0.14 fold compared to control), however this is not statistically significant due to variations in the control samples. Overall, qPCR validation of the disease transcript/FL expression level showed no statistically difference between *ABCA4* sgRNA-treated groups and the control for both disease isoforms (Fig. 4E). In addition, Sanger sequencing confirmed that the 514 bp PCR amplicon corresponded to the *ABCA4* FL transcript containing exons 39 and 40, while the 390 bp amplicon represented *ABCA4* with exon 39 skipping and the 260 bp amplicon represented *ABCA4* with exon 39-40 skipping (Fig. 4F). Altogether, our results showed that RBFOX1N-dPspCas13b-C can reduce mis-spliced variants for *ABCA4* c.5461-10T>C by up to 80%.

### Protein structure prediction of ABCA4 c.4773+3A>G variants with deletion of exon 33-34

To further evaluate the potential of RNA-targeting CRISPR-Cas13 systems in splicing isoform modulation, we tested the designed CRISPR-Cas13 splicing modulating systems for another *ABCA4* variant, c.4773+3A>G which causes skipping of exon 33 or exons 33 and 34. Our *in silico* analysis indicated that skipping of exons 33 and 34 causes a deletion in the ABCA4 exocytoplasmic domain, thus leading to truncated protein shown in the difference between predicted structures of WT ABCA4 and the mutant variants (Fig. 5A-C). Comparison of the verified human ABCA4 protein structure with the mutant proteins also showed misalignment and highlighted the abnormal exocytoplasmic domain encoded by exon 33 and 34 resulting in dysfunctional protein products (Fig. 5D &E).

**Figure 5.**
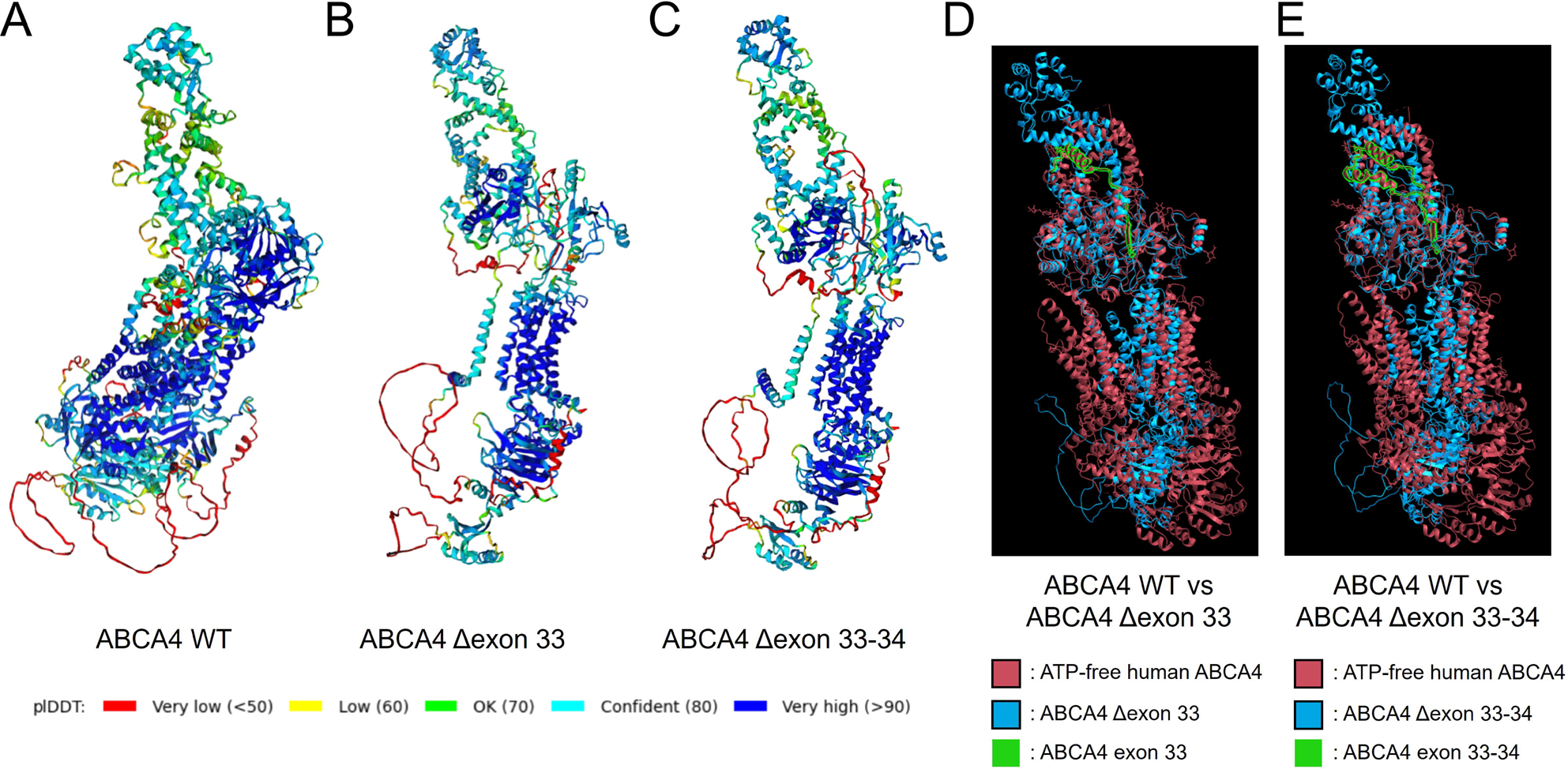
Structural prediction and comparison of the ABCA4 variants with exon 33-34 deletion. (A) Protein structures of the wildtype ABCA4, (B) the ABCA4 Δexon 33, and (C) the ABCA4 Δexon 33-34 predicted by the ColabFold tool. The amino acid residues were coloured based on the plDDT score, with a higher score representing higher confidence in the accuracy of the predicted structure. Regions below 50 plDDT may be unstructured in isolation. (D) Comparison of protein structures of ABCA4 Δexon 33 and WT using ChimeraX. The verified ATP-free human ABCA4 protein structure was used as a reference (red, ^27^). ABCA4 Δexon 33 was presented in blue. The peptide encoded by exon 33 was highlighted in green. (E) Comparison of protein structures of ABCA4 Δexon 33-34 and WT using ChimeraX. The verified ATP-free human ABCA4 protein structure was used as a reference and highlighted in red. ABCA4 Δexon 33-34 was presented in blue. The peptide encoded by exon 33-34 was highlighted in green.

### Evaluation of CASFx to modulate splicing defects of ABCA4 c.4773+3A>G

To test the efficiency of RNA-targeting CRISPR systems for alternative splicing modulation for the *ABCA4* c.4773+3A>G variant, we utilized the *ABCA4* minigene system containing exon 33 and 34 along with the c.4773+3A>G variant (Fig. 6A). In ARPE19, the mutant *ABCA4* minigene showed three distinctive splicing products; a FL product containing both exon 33 and 34 (441 bp), a truncated disease isoform with exon 33 deletion (Δ exon 33, 335 bp), as well as a truncated disease isoform with exon 33 and 34 deletion (Δ exon 33-34, 260 bp) (Fig. 6B & C). First, we tested the potential of the CASFx system to reduce disease isoforms caused by the *ABCA4* c.4773+3A>G variant. Two sgRNAs were designed to target the intron downstream of exon 33 (Fig. 6A). Using the CASFx-1 system, treatment with sgRNA 1, sgRNA 2 or sgRNA1+2 combination showed no significant effect to reduce the Δ exon 33 or Δ exon 33-34 disease isoforms (sgRNA1+2: Δ exon 33 = 0.93 fold, Δ exon 33-34 = 0.87 fold compared to control, Fig. 6C & D). Similar results were also observed in the CASFx-3 system (sgRNA1+2: Δ exon 33 = 1.05 fold, Δ exon 33-34 = 0.93 fold compared to control, Fig. 6C & D). Further quantification of the disease/FL transcript ratio also showed no significant difference in the CASFx-treated groups compared to MT (Fig. 6E). In summary, we did not observe a significant effect of CASFx-1 and CASFx-3 in modulating the level of splice variants of *ABCA4* c.4773+3A>G in ARPE19.

**Figure 6.**
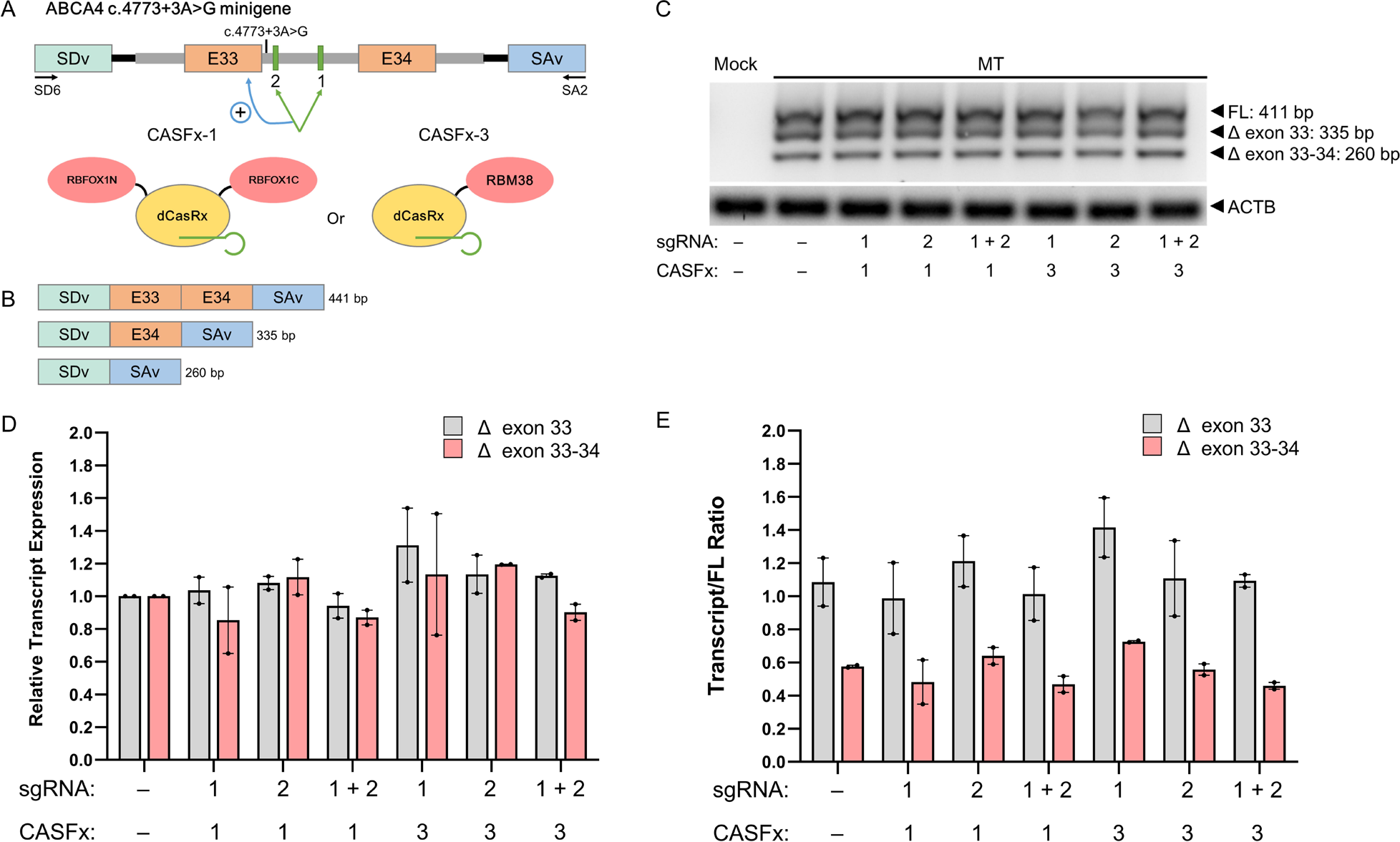
Evaluation of CASFx to modulate splicing defects of ABCA4 c.4773+3A>G. (A) A schematic illustration of the pSPL3–*ABCA4* c.4773+3A>G minigene and the CRISPR splicing factors CASFx-1 (RBFOX1N-dCasRx-C) and CASFx-3 (dCasRx-RBM38), as well as two sgRNA target sites downstream of exon 33 (E33) (green). Positions of SD6 and SA2 primers for splicing product amplification are displayed. (B) Expected splicing products from the *ABCA4* c.4773+3A>G minigene. Normal splicing of the minigene gives rise to a FL product containing both exon 33 and 34 (441 bp) while skipping of exon 33 or exon 33 and 34 generates shorter isoforms (335 and 260 bp). (C) RT-PCR results of ABCA4 splice variants in ARPE19 cells transfected with the ABCA4 minigene system (MT), CASFx-1, CASFx-3 and specified sgRNAs. *ACTB* was used as a housekeeping gene. Mock control represents ARPE19 without transfection of the ABCA4 minigene system. (D) Quantification of relative transcript ratio of *ABCA4* splice variants and (E) ratio of disease transcript to FL transcript compared to MT control without treatment. Data are represented as mean ± SEM (n = 2).

### Evaluation of dPspCas13b to modulate splicing defects of ABCA4 c.4773+3A>G

We next assessed the effect of RBFOX1N-dPspCas13b-C to reduce the disease isoforms caused by *ABCA4* c.4773+3A>G. Six sgRNAs were designed targeting the c.4773+3A>G variant downstream of exon 33 (Fig. 7A). Notably we observed a significant effect of RBFOX1N-dPspCas13b-C in reducing the *ABCA4* disease isoforms. Among all six sgRNAs tested in the ARPE19 cells, sgRNA4 and sgRNA5 showed the strongest effect in reducing the Δ exon 33 and Δ exon 33-34 disease isoform, while sgRNA3 had a limited effect on reducing Δ exon 33-34, while sgRNA1, 2 and 6 had no significant effect (Fig. 7B). Further quantification confirmed the significant effect of sgRNA4 and sgRNA5 in reducing both Δ exon 33 (51.3% and 51.9% reduction respectively), and the Δ exon 33-34 (48.7% and 55.1% respectively) disease isoforms, as well as sgRNA3 in lowering the Δ exon 33-34 isoform (37.2% reduction, Fig. 7C).

**Figure 7.**
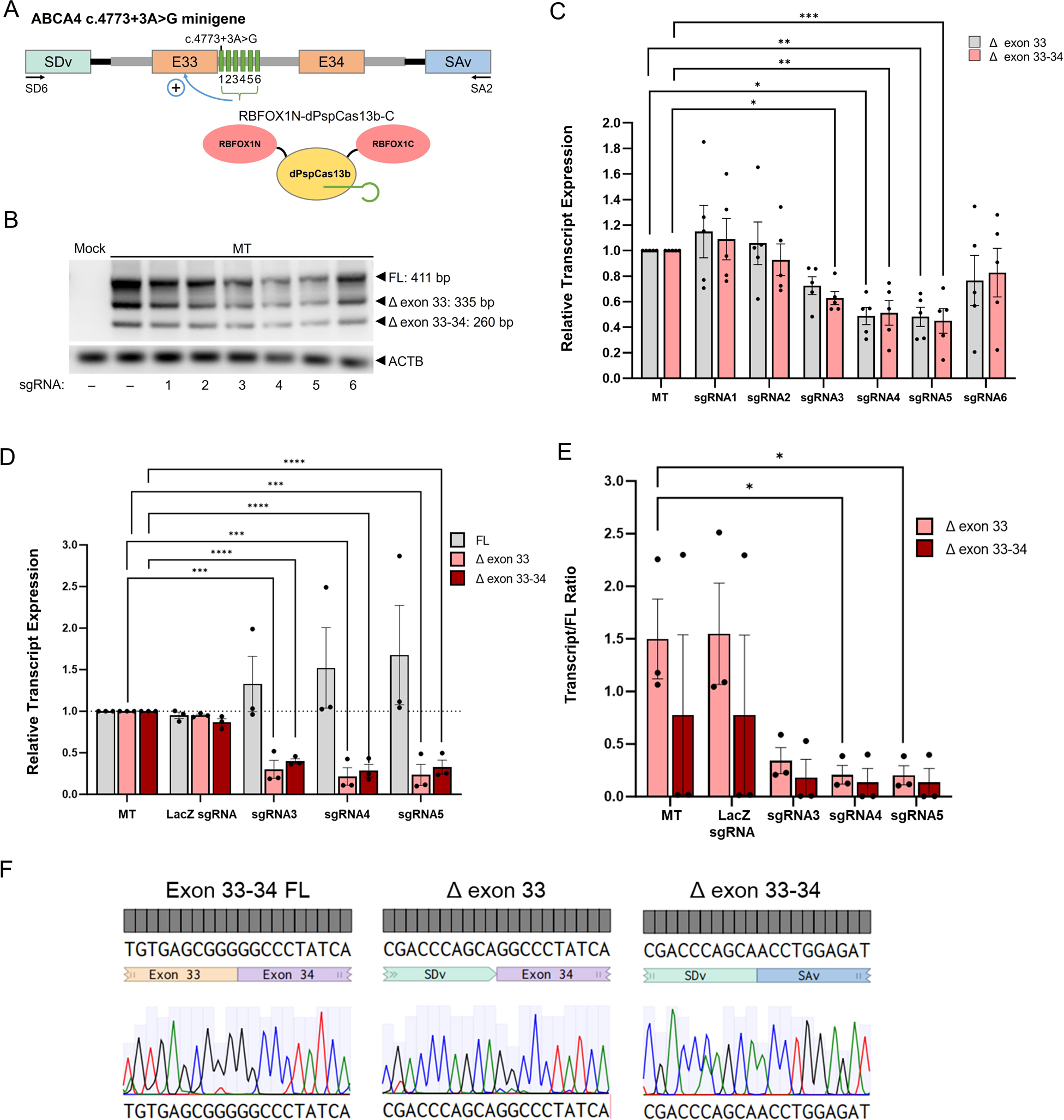
Evaluation of dPspCas13b to modulate splicing defects of ABCA4 c.4773+3A>G. (A) A schematic illustration of the pSPL3–ABCA4 c.4773+3A>G minigene and the CRISPR splicing factor RBFOX1N-dPspCas13b-C, as well as six sgRNA target sites downstream of E33 (green). Positions of SD6 and SA2 primers for splicing product amplification are displayed. (B) RT-PCR results of ABCA4 splice variants in ARPE19 cells transfected with the ABCA4 minigene system (MT), RBFOX1N-dPspCas13b-C and specified sgRNAs. ACTB was used as a housekeeping gene. Mock control represents ARPE19 without transfection of the ABCA4 minigene system. (C) Quantification of relative transcript ratio of ABCA4 splice variants. Data are represented as mean ± SEM (n = 5). *: p < 0.05, **: p < 0.01, ***: p < 0.005. (D) qPCR quantification of the relative expression level of ABCA4 splice variants and (E) relative expression level of disease transcript to FL transcript compared to MT control without treatment. Data are presented as mean ± SEM (n = 3). *: p < 0.05, ***: p < 0.005, ****: p < 0.0001. (F) Sanger sequencing confirmed the presence of ABCA4 FL transcript (left), the Δexon 33 isoform (middle) and the Δexon 33-34 isoform (right).

We then performed qPCR validation for selected sgRNA. Our results showed significant decrease in both Δ exon 33 and Δ exon 33-34 isoforms treated by sgRNA3 (69.8% and 60.16% reduction respectively), sgRNA4 (78.4% and 71.5% reduction respectively) and sgRNA5 (76.5% and 67.1% respectively, Fig. 7D). In addition, LacZ sgRNA showed no effect on expression levels of the FL and disease isoforms. Further quantification of the disease/FL transcript ratio showed a significant decrease in the Δ exon 33/FL transcript ratio treated with sgRNA4 (86.1% reduction) and sgRNA5 (86.5% reduction) compared to control. There was also a sharp decrease in Δ exon 33-34/FL transcript ratio treated with sgRNA4 and sgRNA5 (82.5% and 82.7% reduction compared to control), although not statistically significant due to sample variations (Fig. 7E). Both disease/FL transcript ratios were not affected by LacZ sgRNA treatment. Finally, Sanger sequencing confirmed that the 441 bp amplicon corresponded to *ABCA4* FL transcript containing exons 33 and 34, the 335 bp amplicon represented a truncated *ABCA4* with exon 33 skipping and the 260 bp amplicon represented *ABCA4* transcript with exon 33-34 skipping (Fig. 7F). Altogether, our results showed that RBFOX1N-dPspCas13b-C can reduce the mis-spliced variant of *ABCA4* c.5461-10T>C by up to 78%.

### Evaluation of engineered U1s to modulate splicing defects of ABCA4 c.5461-10T>C and c.4773+3A>G

Beyond the CRISPR-Cas13 systems, we also explored an alternative method for modulating alternative splicing using ExSpeU1s. For *ABCA4* c.5461-10T>C mutation, we designed 4 ExSpeU1s targeting the mutation site (Fig. 8A). The ExSpeU1s were packaged into lentivirus and tested in ARPE19 cells transfected with the *ABCA4* minigene system. Overall, our RT-PCR results showed no significant changes to both disease transcript levels and disease/FL transcript ratios in any of the ExSpeU1 treatments (snRNA4: Δ exon 39 = 0.87 fold, Δ exon 39-40 = 0.98 fold, Δ E39/FL = 1.02 fold, Δ E39-40/FL = 1.08 fold compared to control, Fig. 8B-D).

**Figure 8.**
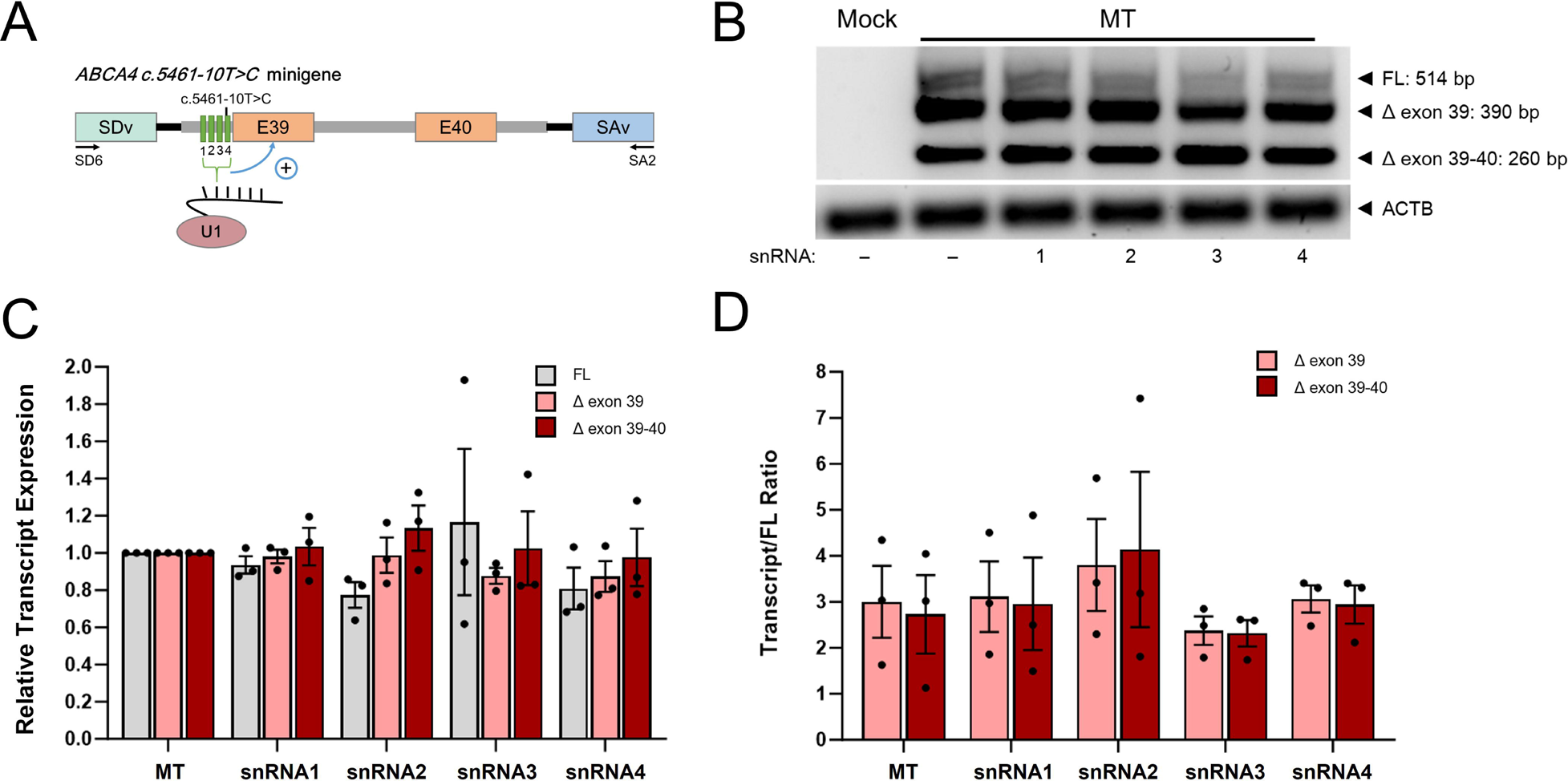
Evaluation of ExSpeU1 to modulate splicing defects of *ABCA4 c.5461-10T>C* mutation. (A) A schematic illustration of the pSPL3–ABCA4 c.5461-10T>C minigene and 4 engineered ExSpeU1 snRNA target sites upstream of E39 (green). Positions of SD6 and SA2 primers for splicing product amplification are displayed. (B) RT-PCR results of *ABCA4* splice variants in ARPE19 cells expressing the *ABCA4* minigene system (MT) and the ExSpeU1s. *ACTB* was used as a housekeeping gene. Mock control represents ARPE19 without transfection of the *ABCA4* minigene system. (C) Quantification of relative transcript ratio of *ABCA4* splice variants and (D) ratio of disease transcript to FL transcript. Data are represented as mean ± SEM (n = 3).

We next tested the efficacy of ExSpeU1 as an alternative splicing modulator in the *ABCA4* c.4773+3A>G mutation. Similarly, 6 ExSpeU1s were designed targeting the mutation site (Fig. 9A). RT-PCR analysis revealed that snRNA2 significantly reduced both Δ exon 33 and Δ exon 33-34 transcript levels (27% and 50% reduction respectively, Fig. 9B-9C). Interestingly, snRNA2 also significantly elevated the level of FL transcripts (1.86 fold increase, Fig. 9C). Further validation with qPCR analysis also confirmed a significant decrease in expression levels of both disease isoforms treated with snRNA2 (Δ exon 33 = 52.1% reduction; Δ exon 33-34 = 84% reduction, Fig. 9D). Additionally, the disease/FL transcript ratios for both disease isoforms were reduced by the snRNA2 treatment (ΔE33/FL = 0.36 fold; ΔE33-34/FL = 0.1 fold compared to control), indicating a reduction in the disease isoforms while the FL transcript was maintained (Fig. 9E). Altogether, these results highlighted the capability of ExSpeU1 to reduce *ABCA4* c.4773+3A>G disease variants by up to 84% and shift the alternative splicing pattern.

**Figure 9.**
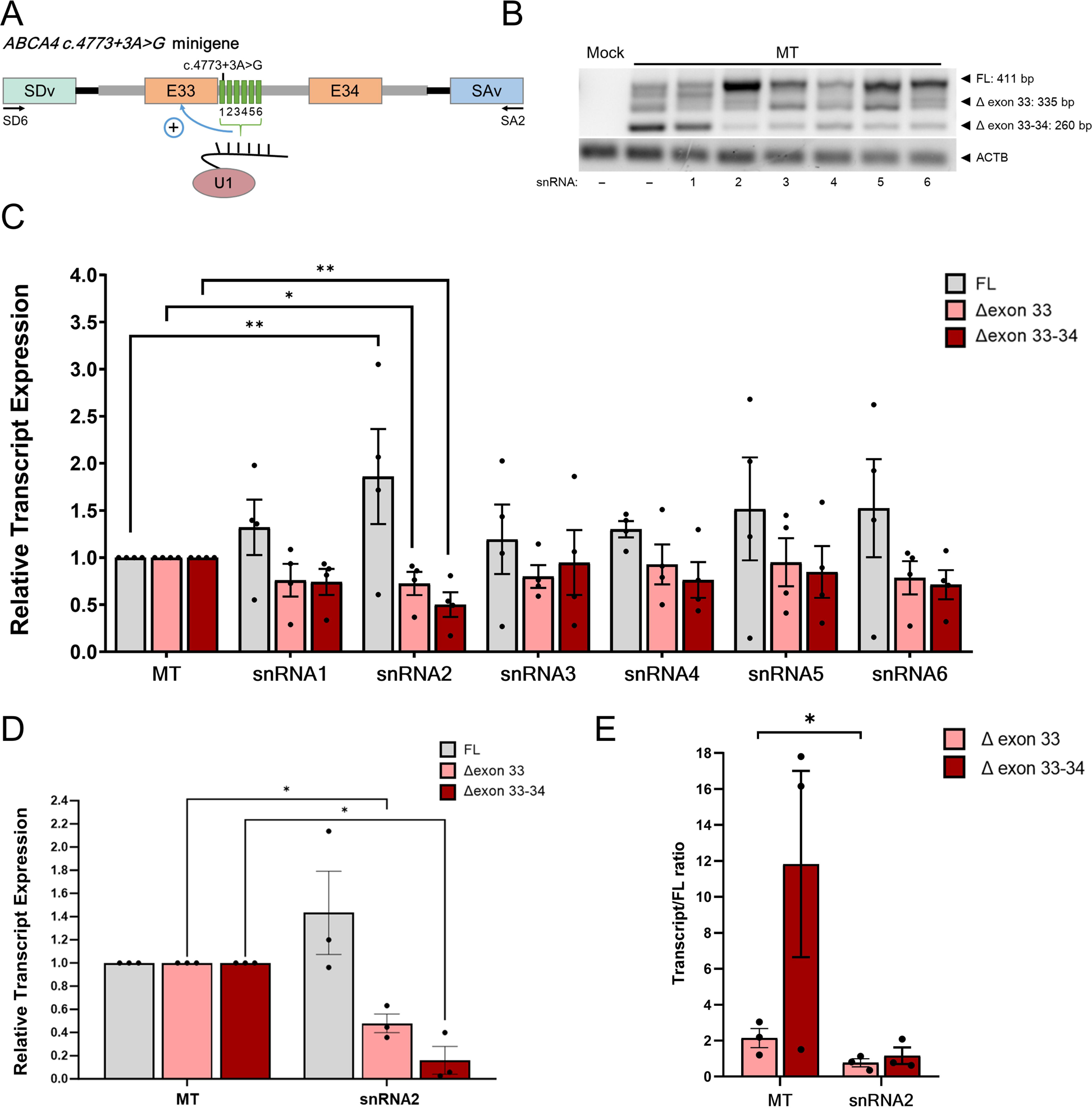
Evaluation of ExSpeU1 to modulate splicing defects of *ABCA4 c.4773+3A>G* mutation. (A) A schematic illustration of the pSPL3–*ABCA4* c.4773+3A>G minigene and 6 engineered ExSpeU1 target sites downstream of E33 (green). Positions of SD6 and SA2 primers for splicing product amplification are displayed. (B) RT-PCR results of *ABCA4* splice variants in ARPE19 cells expressing the *ABCA4* minigene system (MT) and the ExSpeU1s. *ACTB* was used as a housekeeping gene. Mock control represents ARPE19 without transfection of the *ABCA4* minigene system. (C) Quantification of relative transcript expression of *ABCA4* splice variants. Data are represented as mean ± SEM (n = 3). *: p < 0.05, **: p < 0.01. (D) qPCR quantification of the relative expression level of *ABCA4* splice variants and (E) relative expression level of disease transcript to FL transcript compared to MT control without treatment. Data are presented as mean ± SEM (n = 3). *: p < 0.05.

## Discussion

The dysregulation of the alternative splicing process is one of the contributors to STGD1, as well as a broad range of diseases including cardiomyopathies, hereditary peripheral neuropathies, muscular dystrophies and premature aging syndromes ^28–37^. Various therapeutic strategies have been developed to target the splicing mechanism to correct these mis-splicing events, such as antisense oligonucleotides (AONs), as well as small molecules that regulate splicing enhancers and silencers ^38^. Notably, a recent report demonstrated the use of AON to restore splicing defects for *ABCA4* c.5461-10T>C ^39^. Such strategies using AON have gained success, leading to FDA-approved treatments for Duchenne Muscular Dystrophy ^40,41^ and spinal muscular dystrophy ^42^. To expand the molecular toolset for modulating splicing defects, in this study we tested the use of RNA-targeting CRISPR-Cas13 and engineered U1 systems. Table 1 summarized the performance of the RNA-targeting CRISPR-Cas13 and U1 systems tested in this study to modulate splicing defects of *ABCA4* c.5461-10T>C and c.4773+3A>G. By application of RNA-targeting CRISPR systems and specific sgRNAs, we were able to reduce the level of *ABCA4* mis-spliced transcripts by up to 80%. In comparison, engineered U1 snRNA treatments were able to reduce the level of *ABCA4* mis-spliced transcripts up to 84%.

**Table 1:**
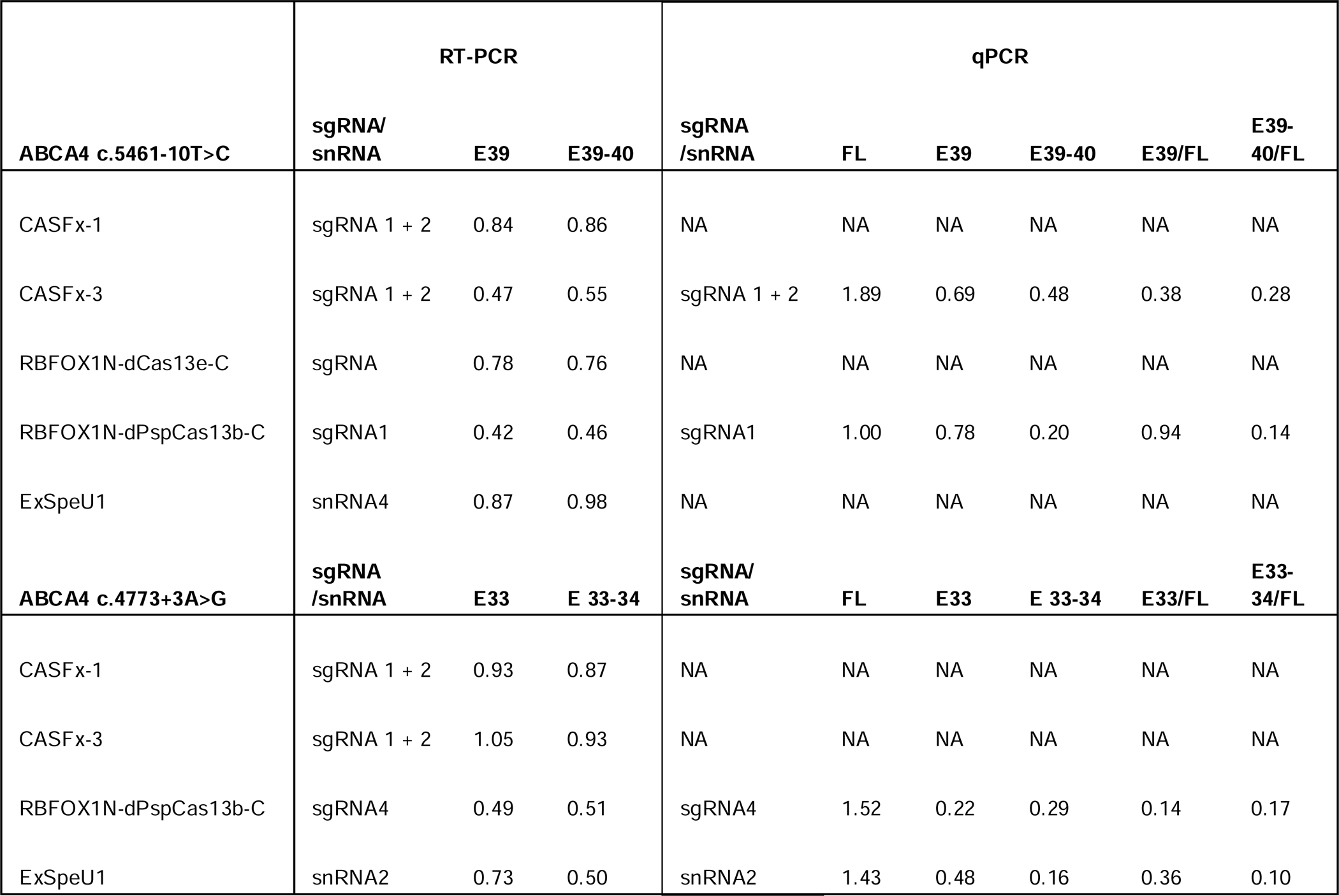
Summary of performance of RNA-targeting CRISPR-Cas and ExSpeU1 systems on modulating ABCA4 splicing defects. The best performing sgRNA/snRNA was listed. For individual transcripts, values indicated mean expression levels normalized to MT control. For diseases transcript to full length ratio, values indicated the ratio normalized to MT control. FL: full length; E39: exon 39; E40: exon 40; E33: exon 33; E34: exon 34; NA: not available.

| ABCA4 c.5461-10T>C | RT-PCR |  |  | qPCR |  |  |  |  |  |
| --- | --- | --- | --- | --- | --- | --- | --- | --- | --- |
|  | sgRNA/<br>snRNA | E39 | E39-40 | sgRNA<br>/snRNA | FL | E39 | E39-40 | E39/FL | E39-40/FL |
| CASFx-1 | sgRNA 1 + 2 | 0.84 | 0.86 | NA | NA | NA | NA | NA | NA |
| CASFx-3 | sgRNA 1 + 2 | 0.47 | 0.55 | sgRNA 1 + 2 | 1.89 | 0.69 | 0.48 | 0.38 | 0.28 |
| RBFOX1N-dCas13e-C | sgRNA | 0.78 | 0.76 | NA | NA | NA | NA | NA | NA |
| RBFOX1N-dPspCas13b-C | sgRNA1 | 0.42 | 0.46 | sgRNA1 | 1.00 | 0.78 | 0.20 | 0.94 | 0.14 |
| ExSpeU1 | snRNA4 | 0.87 | 0.98 | NA | NA | NA | NA | NA | NA |
| ABCA4 c.4773+3A>G | sgRNA<br>/snRNA | E33 | E 33-34 | sgRNA/<br>snRNA | FL | E33 | E 33-34 | E33/FL | E33-34/FL |
| CASFx-1 | sgRNA 1 + 2 | 0.93 | 0.87 | NA | NA | NA | NA | NA | NA |
| CASFx-3 | sgRNA 1 + 2 | 1.05 | 0.93 | NA | NA | NA | NA | NA | NA |
| RBFOX1N-dPspCas13b-C | sgRNA4 | 0.49 | 0.51 | sgRNA4 | 1.52 | 0.22 | 0.29 | 0.14 | 0.17 |
| ExSpeU1 | snRNA2 | 0.73 | 0.50 | snRNA2 | 1.43 | 0.48 | 0.16 | 0.36 | 0.10 |

ABCA4 is a transporter protein located in the disc membrane of the photoreceptor’s outer segments. It is also expressed on the apical membrane of human RPE cells and is responsible for lipid homeostasis ^27^. The experimental structure of the ABCA4 protein has been confirmed using cryogenic electron microscopy previously ^27^. ABCA4 is a single elongated polypeptide consisting of two homologous halves, each containing a transmembrane domain, an exocytoplasmic domain, a cytoplasmic nucleotide-binding domain, and a regulatory domain. In our study, the wild-type ABCA4 protein was accurately predicted as a monomer by ColabFold, albeit still having some structural differences compared to the verified experimental structure. The *ABCA4* exon 39-40 encodes amino acid (aa) 1821-1905, which comprises the exocytoplasmic helices of the second transmembrane domain. Although these structures have not been related to any known function, it is typical for ion channels such as K^+^ or Cl^−^ channels to have helices with one end exposed to the membrane. It is believed that these helices function for ion binding and selection. Skipping of exon 39 or 39-40 caused by the c.5461-10T>C variant results in a premature stop codon at aa. 1832 and aa. 1825 respectively, leading to the formation of a truncated ABCA4 protein without part of the transmembrane domain and the following nucleotide-binding domain and regulatory domain of the second half of the ABCA4 protein. On the other hand, The *ABCA4* exon 33-34 encodes the aa. 1557-1616, which resides within the second exocytoplasmic domain. Both exocytoplasmic domains have not been reported to have any known enzymatic or structural function, however numerous STGD1 mutations were located in the exocytoplasmic domains, indicating the pivotal role of these domains for ABCA4 functionality. Removal of exon 33 or 33-34 caused by c.4773+3A>G variant leads to a premature stop codon at aa. 1573 and aa. 1600 respectively, resulting in a truncated protein with defective exocytoplasmic and transmembrane domains and deletion of nucleotide-binding domain of the second half of the ABCA4 protein. It is also possible that the premature stop codon in these *ABCA4* variants would trigger nonsense-mediated decay (NMD). Altogether, these ABCA4 defects impact the clearance of toxic retinoid compounds from the photoreceptors and ultimately lead to macular degeneration.

In this study we tested three RNA-targeting CRISPR-Cas13 systems from the Cas13 family for alternative splicing modulation of the *ABCA4* c.5461-10T>C and c.4773+3A>G variants, namely, the CASFxs, Cas13e, and PspCas13b. As shown previously, CASFx-1 has better efficiency in exon inclusion of the *SMN2* gene compared to CASFx-3 ^13^. However, in our study CASFx-1 was unable to modulate *ABCA4* splicing patterns for both c.5461-10T>C and c.4773+3A>G variants. In contrast, CASFx-3 can induce up to 53% reduction of *ABCA4* disease isoforms using a combination of 4 sgRNAs, but CASFx-3 was unable to reduce disease isoforms of the c.4773+3A>G variant. Furthermore, we tethered the alternative splicing factor RBFOX1 to the catalytically inactive dCas13e and built an all-in-one vector with sgRNA targeting the *ABCA4* c.5461-10T>C variant. However, we did not observe exon inclusion using the RBFOX1N-dCas13e-C system. It is possible that further testing with sgRNA targeting different regions may improve the effect of the CASFx or RBFOX1N-dCas13e1-C systems to modulate *ABCA4* splicing. Future research using more efficient delivery method of the CRISPR systems in ARPE19 cells may further improve the effect observed with CASFx or RBFOX1N-dCas13e-C systems. Also, it would be interesting to investigate whether fusing dCas13e with another alternative splicing regulator RBM38 would exert a stronger effect on splicing regulation.

Compared to the CASFx system, RBFOX1N-dPspCas13b-C exhibited greater efficacy in exon inclusion of the *SMN2* gene ^13^. Notably, our results showed that RBFOX1N-dPspCas13b-C indeed exerted the highest efficiency in reducing disease isoforms for both *ABCA4* c.5461-10T>C and c.4773+3A>G compared to all other CRISPR-Cas13 systems tested (50-80% reduction). It is worth noting that for the *ABCA4* c.5461-10T>C variant, dPspCas13b sgRNAs were designed to target the mutation site upstream of exon 39, while the sgRNAs for both CASFxs and dCas13e effectors were targeting the intron downstream of the target exon 39 and 40. Although the general guideline for Cas13 sgRNA design for splicing modulation recommended that the sgRNAs target the splicing donor and acceptor sites of the downstream intron of the target exon, the optimal sgRNA targeting site remain unclear and could varies in different cases. Our study highlighted that it is pivotal to screen through a panel of sgRNA candidates to identify the target site with the best efficiency for each individual case. Notably, the dPspCas13b splicing effector tested in this study is relatively small in size (1465 aa), and can be potentially packaged into AAV, which highlighted its translational value as a retinal gene therapy.

In their bacterial or archaeal origins, RNA-targeting CRISPR-Cas systems serve as “the second line of immunity” and activate with the presence of viral RNA. Given the mutation-prone nature of viral replication and transcription, RNA-targeting effectors have developed a more relaxed targeting specificity, having higher tolerance of mismatches within recognition sequences ^43,44^. This leads to a higher off-target activity when RNA-targeting CRISPR-Cas systems are put into therapeutic application ^45^. In our case of CRISPR-based alternative splicing modulation strategy, the consequences of off-targets would be less severe as we are using catalytically inactive forms of CRISPR endonuclease proteins which act on RNA, making it unlikely to cause permanent deleterious genetic modifications. Additionally, the CRISPR systems need to target specific sites where the splicing machinery exerts its function. In support of this, we did not observe any off-target effects with LacZ sgRNA treatment by qPCR analysis. As such, RNA-targeting CRISPR-Cas13 systems provide a promising approach for transcriptome engineering.

On the other hand, U1 snRNA is part of the spliceosomal machinery responsible for catalyzing the alternative splicing process. The spliceosome is composed of snRNAs and five major small nuclear ribonucleoproteins (snRNPs) named U1, U2, U4, U5, and U6. The splicing process begins with the recognition of the 5’ splice site by U1 snRNP at the 5’ exon-intron junction and the binding of U2 snRNP at the branch point sequence within the intron. The other snRNPs then join and the spliceosome undergoes a series of structural changes, forming a catalytically active spliceosome. The active spliceosome excises introns and forms a lariat structure, while ligating the adjacent exons together, resulting in the formation of a mature mRNA ^46^. U1 snRNPs recognise the 5’ splice site through base-pairing interactions between the 10 highly conserved nucleotides at the 5’ end of U1 snRNAs and the intron sequences of the pre-mRNA. When alternative splicing occurs, different splice sites within the pre-mRNA may compete for recognition by the spliceosome. This highlights the pivotal role of U1 snRNAs for the initiation and selection of the 5’ splice sites, making it applicable to determine whether an exon is included or excluded in the mature mRNA by specifying the desired U1 snRNA sequences.

As U1 snRNA determines the 5’ splice site of the alternative splicing process, the designed ExSpeU1 sequences are required to target the downstream intronic region of the target exon to facilitate exon inclusion ^22^. In our testing, ExSpeU1 treatment induced the inclusion of exon 33 and 34 when targeting the *ABCA4* c.4773+3A>G mutation downstream of the target exon, showing a decrease in disease mis-spliced transcripts and elevation in FL transcript. However, ExSpeU1 treatment exerted no effect for *ABCA4* c.5461-10T>C mutation as the ExSpeU1s were targeting the mutation upstream of the target exon. It is possible that further testing with ExSpeU1 targeting downstream of exon 39 may improve the effect of the ExSpeU1 to correct the *ABCA4* c.5461-10T>C splicing defect. Nevertheless, our results demonstrated great potential in modulating the alternative splicing pattern by delivery of a single ExSpeU1 component. Additionally, the U1 snRNA is typically 164 bases long in humans, making it an appropriate target to be packaged into AAVs for clinical applications. To our knowledge, our work represents the first proof-of-concept study to demonstrate the use of engineered U1 to modulate *ABCA4* splicing defects.

In summary, our study illustrated the feasibility of employing RNA-targeting CRISPR-Cas technology and ExSpeU1 to mitigate *ABCA4* disease isoforms, providing a novel approach to treat STGD1 caused by *ABCA4* mis-splicing. By delivering specified gRNAs and dPspCas13b splicing effectors, we have successfully attenuated the levels of *ABCA4* disease isoforms associated with the c.5461-10T>C and c.4773+3A>G splicing mutations. Additionally, delivery of a single ExSpeU1 has proven effective in diminishing disease isoforms arising from the *ABCA4* c.4773+3A>G mutation. By reducing the presence of disease isoforms, we potentially mitigate the adverse effects of the mis-splicing mutations, underscoring the significant promise of RNA-targeting CRISPR-Cas systems and ExSpeU1 for clinical applications in STGD1 treatment.

## Materials and Methods

### sgRNA design and plasmid construction

Supplementary table 1 summarized the sgRNAs used in this study. Two *ABCA4* minigene constructs carrying the variants c.4773+3A>G and c.5461-10T>C respectively were constructed previously ^7^. CASFx-1 (RBFOX1N-dCasRx-C) and CASFx-3 (dCasRx-RBM38) were obtained from Addgene (Plasmid #118635 and #118638). RBFOX1N-dPspCas13b-C was constructed previously ^13^. For CASFx, sgRNA were designed based on guidelines previously reported ^47,48^. sgRNAs are designed to target 5 bp downstream of the splicing donor site and the middle of the intron for each target exon. We synthesized sgRNA expression cassettes containing the sequences of a U6 promoter, the CasRX direct repeat, followed by the specific sgRNA. gBlocks were ordered from IDT and amplified with High Fidelity PCR reaction using the following primers: Forward: 5’-TGAGTATTACGGCATGTGAGGGC-3’, Reverse: 5’-TCAATGTATCTTATCATGTCTGCTCGA-3’. PCR amplification of sgRNA expression cassettes was performed as described previously ^49,50^. A region of the LacZ sequence was used as a non-targeting control sgRNA sequence (TGCGAATACGCCCACGCGAT) for CASFxs and RBFOX1N-dPspCas13b-C as it does not target any gene in mammalian cells.

For the construction of RBFOX1N-dCas13e-C, Cas13e protein sequence was a gift from Dr Hui Yang ^15^. The ABCA4 sgRNA sequence was from Dr Fred Chen’s group based on the sequence of an AON: CAGGGCCCATGCTCCATGGGCCTCG. A region of the mCherry sequence was used as a non-targeting control sgRNA sequence (TCCTCGAAGTTCATCACGCGCTCCCACTTGAAGCCCTCGGGGAAGGACAG) as it does not target any gene in mammalian cells. Cas13e was truncated from the C-terminal to remove the HEPN2 domain, resulting in a truncated Cas13e protein of 625 aa. Dead Cas13e (dCas13e) was obtained through site-directed mutagenesis (R84A and H89A). RBFOX1N was fused to the N and C-terminal of truncated dCas13e to produce RBFOX1N-dCas13e-C cassette. RBFOX1N-dCas13e-C cassette was then cloned into pAAV-MCS vector after CMV promoter to produce CMV-RBFOX1N-dCas13e-C. The U6 promoter-driven gRNA sequences (ABCA4 or mCherry) were then cloned into the pCMV-RBFOX1N-dCas13e-C vector to generate the all-in-one U6-gRNA-CMV-RBFOX1N-dCas13e-C constructs (Addgene #206049 and 206050).

### Transfection of CRISPR-Cas RNA splicing modulating systems into ARPE19 cells

For transfection, 3×10^4^ ARPE19 cells were seeded in 12-well plates on Day 0. Two transfections were carried out using Lipofectamine 3000 (Thermo Fisher Scientific) according to the manufacturer’s instructions. On Day 1, the cells were first transfected with 800 ng of minigene constructs, followed by transfection with CRISPR and sgRNA plasmids on Day 2. For CASFx, 800 ng of CRISPR plasmids and 360 ng of gBlock are used for transfection. For RBFOX1N-dPspCas13b-C and RBFOX1N-dCas13e-C, 1000 ng of CRISPR and sgRNA plasmids are used for transfection. Cells were harvested for RNA extraction on Day 6.

### ExSpeU1 construction and lentivirus generation

8 ExSpeU1 snRNAs were cloned into the LentiGuide vector backbone (Supplementary table 2). Lentivirus was generated using the third-generation packaging system by transfecting the lentiguide-ExSpeU1 snRNA lentiviral vectors together with the packaging plasmids pMDLg/pRRE, pRSV-Rev, and pMD2.G (all from Addgene) into HEK293FT cells using PEI transfection reagent. 5×10^6^ HEK293FT cells were plated on a 10 cm^2^ tissue culture dish the day before transfection. The supernatant containing the lentivirus was collected 48 and 72 hours after transfection. The pooled supernatant was then filtered with 0.45 μm membrane and concentrated using the PEG-it virus precipitation solution (System Biosciences #LV810A-1) according to the manufacturer’s instructions. Lentivirus titration was determined using Lenti-X p24 Rapid Titre Kit (Takara #631476) according to the manufacturer’s instructions. Lentivirus was aliquoted and stored at −80°C.

### Lentiviral transduction of ExSpeU1 into ARPE19 cells

Human retinal pigment epithelium ARPE19 cells were cultured in DMEM medium with 10% fetal bovine serum, 1X Penicillin-Streptomycin, and 1X Glutamax (all from Thermo Fisher Scientific), using an incubator at 37°C and 5% CO2. For lentiviral transduction of ExSpeU1 to modulate *ABCA4* alternative splicing, 3×10^4^ ARPE19 cells were seeded in 12-well plates on Day 0. Transfection of 800 ng of minigene constructs was first carried out on Day 1 using Lipofectamine 3000 (Thermo Fisher Scientific) according to the manufacturer’s instructions. On Day 2 the media were replaced and the cells were transduced with U1 snRNA lentivirus by adding the lentivirus at a multiplicity of infection (MOI) of 30 directly to the cells along with 8 µg/mL polybrene. The media were changed into fresh media the day after transduction. Cells were harvested for RNA extraction on Day 6.

### RT-PCR assay for ABCA4 splice variants detection

Total RNA from cell samples was extracted using the Illustra RNAspin kit with DNaseI treatment (GE Healthcare). RT-PCR was performed with 100 ng of total RNA using SuperScript III One-Step RT-PCR System (Thermo Fisher Scientific) according to the manufacturer’s instructions. The RT-PCR thermal profile used was 55°C for 30 min then 94°C for 2 min for cDNA synthesis, followed by 30 cycles of the PCR process: 30 s of 94°C (denaturation), 1 min of 56.5°C (annealing), and 2 min of 68°C (extension). The RT-PCR products were amplified using the pSPL3-specific primers SD6: 5’-TCTGAGTCACCTGGACAACC-3’ and SA2: 5’-ATCTCAGTGGTATTTGTGAGC-3’. The RT-PCR products were then visualized on 1.5% agarose gel. ACTB was used as a housekeeping gene. RT-PCR for ACTB was performed using the forward primer 5’-CCCTGGCACCCAGCAC-3’ and reverse primer 5’-GCCGATCCACACGGAGTAC-3’, following the same RT-PCR condition described above except with 61.9°C for annealing temperature.

### qPCR assay for ABCA4 splice variants detection

Total RNA from cell samples was extracted and subjected to DNaseI treatment using the Illustra RNAspin kit (GE Healthcare). cDNA synthesis from the RNA samples was performed using the High-Capacity cDNA Reverse Transcription Kit with RNase Inhibitor (Applied Biosystems #4374966) according to the manufacturer’s instruction. Quantitative PCR was then carried out with the PowerUp SYBR Green Master Mix (Applied Biosystems #A25742) with specific primers targeting individual *ABCA4* transcripts (Supplementary Table 3). The subsequent qPCR analysis was performed with the StepOnePlus™ Real-Time PCR System and Software (Applied Biosystems) using an initial denaturation at 95°C for 2 min followed by 40 cycles of PCR stage at 60°C for 30 sec. A melting curve analysis was also performed to confirm a single PCR product.

### Statistical analysis

Statistical analysis was performed using paired t-test or one-way ANOVA, followed by Dunnett’s multiple comparison tests where appropriate (Graphpad Prism). Statistical differences were considered significant when P ≤ 0.05.

### Protein structure prediction and comparison using ColabFold and ChimeraX

The protein structures of the WT and mutated ABCA4 proteins were predicted using the ColabFold (ColabFold v1.5.2-patch: AlphaFold2 using MMseqs2) tool ^26^. The prediction was run using mmseq2_uniref_env with unpaired_paired mode. Three recycles were used for each model. In total, five models were generated and ranked based on the Predicted Local Distance Difference Test score (plDDT). The top-ranked models were selected for this study. Structural comparison of WT and mutant ABCA4 proteins was carried out using the ChimeraX tool ^51,52^ with the Matchmaker function. The experimental structure of ATP-free human ABCA4 protein has been published previously ^27^ and was used as a WT reference.

## Competing interests

The authors declare no conflict of interest.

## Author contribution

RCBW and AC designed the experiments; RHCL, DU, SK and FTH conducted the experiments; IMW, IG, GSL, FC, SM and AC provided reagents; RHCL, DU, KM, TE and RCBW analysed the data; KM, TE and RCBW provided funding to this work; RHCL and RCBW wrote the manuscript. All authors approved the manuscript.

## Acknowledgement

This research was funded by the University of Melbourne (RCBW) and the Centre for Eye Research Australia (RCBW). RCBW is supported by the University of Melbourne, the Centre for Eye Research Australia, National Health and Medical Research Council (GCT1184076) and Medical Future Research Fund (MRF2024365). RHCL and DU are supported by the Melbourne Research Scholarship. SK is supported by the Melbourne Research Scholarship and Retina Australia. The Centre for Eye Research Australia receives operational infrastructure support from the Victorian Government. FKC receives support from the WA Health Near-miss Award and the Telethon Grant 2023. FTH is supported by Ministry of Science and Technology, Taiwan, ADD-ON project (ID: MOST108-2314-B-039-007-MY3).

## Notes

### Competing Interest Statement

The authors have declared no competing interest.

