## Supplementary table 1-3 for "Using RNA-targeting CRISPR-Cas13 and engineered U1 systems to reduce *ABCA4* splice variants in Stargardt disease"

**Supplementary table 1:** Information of sgRNAs used in this study.

| **Name** | **Distance from mutation** | **Strand** | **Sequence** |
| --- | --- | --- | --- |
| CASFx exon 33 sgRNA1 | c.4773+3A>G +51 bp | + | GAATGAGAAACTCTCATGAGTGA |
| CASFx exon 33 sgRNA2 | c.4773+3A>G +4 bp | + | AACAGACTGGAGATTTGAGTAGG |
| dPspCas13b exon 33 sgRNA1 | c.4773+3A>G 0 bp | - | CAAATCTCCAGTCTGTTTACAc |
| dPspCas13b exon 33 sgRNA2 | c.4773+3A>G +9 bp | - | AAATCCTACTCAAATCTCCAGT |
| dPspCas13b exon 33 sgRNA3 | c.4773+3A>G +18 bp | - | GCAAGTCAAAAATCCTACTCAA |
| dPspCas13b exon 33 sgRNA4 | c.4773+3A>G +29 bp | - | ATGGTAGTTAAGCAAGTCAAAA |
| dPspCas13b exon 33 sgRNA5 | c.4773+3A>G +40 bp | - | GTTTCTCATTCATGGTAGTTAA |
| dPspCas13b exon 33 sgRNA6 | c.4773+3A>G +49 bp | - | CTCATGAGAGTTTCTCATTCAT |
| CASFx exon 39 sgRNA1 | c.5461-10T>C +287 bp | + | GCCTCTGTTTCCATGGCTGGGGA |
| CASFx exon 39 sgRNA2 | c.5461-10T>C +140 bp | + | GTAGCCGAGGCCCATGGAGCATG |
| CASFx exon 40 sgRNA1 | c.5461-10T>C +1547 bp | + | TGGCCATACCTTTTAGAGGCTTT |
| CASFx exon 40 sgRNA2 | c.5461-10T>C +602 bp | + | CATGCCACACCCTGGGCCAGTGG |
| dCas13e sgRNA (AON) | c.5461-10T>C +145 bp | - | CAGGGCCCATGCTCCATGGGCCTCG |
| dPspCas13b exon 39 sgRNA1 | c.5461-10T>C +4 bp | - | ACAGcGAAGTAGGACTGTTGGAAACGGGGC |
| dPspCas13b exon 39 sgRNA1 | c.5461-10T>C +6 bp | - | AAACAGcGAAGTAGGACTGTTGGAAACGGG |

**Supplementary table 2:** Information of ExSpeU1s used in this study.

| **ExSpeU1** **name** | **Strand** | **snRNA sequence** |
| --- | --- | --- |
| ABCA4 c.4773+3A>G ExSpeU1 snRNA1 | - | TCTCCAGTCTGTTTACAC |
| ABCA4 c.4773+3A>G ExSpeU1 snRNA2 | - | CAAATCTCCAGTCTGTTT |
| ABCA4 c.4773+3A>G ExSpeU1 snRNA3 | - | TACTCAAATCTCCAGTCT |
| ABCA4 c.4773+3A>G ExSpeU1 snRNA4 | - | ATCCTACTCAAATCTCCA |
| ABCA4 c.4773+3A>G ExSpeU1 snRNA5 | - | AAAAATCCTACTCAAATC |
| ABCA4 c.4773+3A>G ExSpeU1 snRNA6 | - | AGTCAAAAATCCTACTCA |
| ABCA4 c.5461-10T>C ExSpeU1 snRNA1 | - | GAAACAGGGAAGTAGGAC |
| ABCA4 c.5461-10T>C ExSpeU1 snRNA2 | - | AACAGGGAAGTAGGACTG |
| ABCA4 c.5461-10T>C ExSpeU1 snRNA3 | - | CAGGGAAGTAGGACTGTT |
| ABCA4 c.5461-10T>C ExSpeU1 snRNA4 | - | GGGAAGTAGGACTGTTGG |

**Supplementary table 3:** Information of optimized qPCR primers used for *ABCA4* transcripts detection.

| **Name** | **Targeting Transcript** | **Sequence** | **Concentration** |
| --- | --- | --- | --- |
| ABCA4 ex33-34-For | ABCA4 c.4773+3A>G FL | 5’-ATCATGAATGTGAGCGGGGG-3’ | 800 nM |
| ABCA4 SAv-Rev-1 | ABCA4 c.4773+3A>G FL  ABCA4 c.4773+3A>G Δexon 33  ABCA4 c.4773+3A>G Δexon 33-34  ABCA4 c.5461-10T>C Δexon 39-40 | 5’-TTCTGATAGGCAGCCTGCAC-3’ | 800 nM |
| ABCA4 SDv-ex34-For | ABCA4 c.4773+3A>G Δexon 33 | 5’-TCTAGAGTCGACCCAGCAGG-3’ | 800 nM |
| ABCA4 SDv-SAv-For | ABCA4 c.4773+3A>G Δexon 33-34  ABCA4 c.5461-10T>C Δexon 39-40 | 5’-GACCCAGCAACCTGGAGATC-3’ | 800 nM |
| ABCA4 e39-e40-For | ABCA4 c.5461-10T>C FL | 5’-GTCTATGCCCGGTTTGGTGA-3’ | 800 nM |
| ABCA4 SAv-Rev-2 | ABCA4 c.5461-10T>C FL  ABCA4 c.5461-10T>C Δexon 39 | 5’-TCAGTGGTATTTGTGAGCCAGG-3’ | 800 nM |
| ABCA4 SDv-e40-For | ABCA4 c.5461-10T>C Δexon 39 | 5’-TAGAGTCGACCCAGCAGTGA-3’ | 800 nM |
| ACTB For | ACTB | 5’-CCCTGGCACCCAGCAC-3’ | 1000 nM |
| ACTB Rev | ACTB | 5’-GCCGATCCACACGGAGTAC-3’ | 1000 nM |
